# *SPOP* mutation confers sensitivity to AR-targeted therapy in prostate cancer by reshaping the androgen-driven chromatin landscape

**DOI:** 10.1101/2021.04.20.440154

**Authors:** Ivana Grbesa, Michael A. Augello, Deli Liu, Dylan R. McNally, Christopher D. Gaffney, Dennis Huang, Kevin Lin, Ramy Goueli, Brian D. Robinson, Francesca Khani, Lesa D. Deonarine, Mirjam Blattner, Olivier Elemento, Elai Davicioni, Andrea Sboner, Christopher E. Barbieri

## Abstract

The normal androgen receptor (AR) cistrome and transcriptional program are fundamentally altered in prostate cancer (PCa). Here, we show that *SPOP* mutations, an early event in prostate tumorigenesis, reshape the chromatin landscape and AR-directed transcriptional program in normal prostate cells. Induction of *SPOP* mutation results in DNA accessibility and AR binding patterns found in human PCa. Consistent with dependency on this AR reprogramming, castration of *SPOP* mutant mouse models results in the loss of neoplastic phenotypes. Finally, human *SPOP* mutant PCa show improved response to AR-targeted therapies. Together, these results show that a single genomic alteration may be sufficient to reprogram the chromatin of normal prostate cells toward oncogenic phenotypes and that *SPOP* mutant tumors may be preferentially dependent on AR signaling through this mechanism.

## Introduction

The androgen receptor (AR) is a critical driver and key therapeutic target in prostate cancer (PCa). In normal prostate epithelial cells, AR coordinates growth-suppressive effects and differentiation programs (Heinlein and Chang, 2004; Taplin and Balk, 2004). During prostate carcinogenesis, AR is reprogrammed to instead promote oncogenic transcriptional programs (Pomerantz et al., 2015). However, the mechanisms of this reprogramming remain incompletely understood.

Recurrent missense mutations in *SPOP* (Speckle Type BTB/POZ Protein) occur in about 10% of localized primary PCa cases (Cancer Genome Atlas Research, 2015; Li et al., 2020) and are highly prostate-specific; they are rarely observed in other cancer types. *SPOP* mutations nominate a distinct molecular subtype of human PCa (Barbieri et al., 2012; Cancer Genome Atlas Research, 2015; Shoag et al., 2018), with defined genomic alterations, DNA methylation, transcriptional signatures, and clinical characteristics (Barbieri et al., 2012; Boysen et al., 2015; Cancer Genome Atlas Research, 2015; Liu et al., 2018). Furthermore, a variety of data suggest that *SPOP* mutations occur early in the process of prostate tumorigenesis (Baca et al., 2013; Boysen et al., 2015). SPOP acts as the substrate recognition component of a CUL3-E3 ubiquitin-protein ligase complex (Zhuang et al., 2009), with prostate cancer-associated mutations affecting substrate specificity. Substrates reported to be affected by *SPOP* mutations include AR itself (An et al., 2014; Geng et al., 2014), the AR coactivators SRC3 (Geng et al., 2013), p300 (Blattner et al., 2017), and TRIM24 (Blattner et al., 2017), and chromatin-associated proteins like DEK (Theurillat et al., 2014) and DAXX (Bouchard et al., 2018). However, how mutations in *SPOP* impact AR-directed transcriptional programs, and if this is required for tumor initiation and progression, remains unclear.

Here we leverage the unique capabilities of 3D prostate organoids to define the epigenomic and transcriptional landscape upon AR activation and utilize a genetically engineered mouse (GEM) model conditionally expressing mutant *SPOP* (Blattner et al., 2017) to characterize the impact of a single genomic alteration in genetically normal cells.

## Results

### Chromatin changes in normal prostate organoids with androgen stimulation

Compared to our understanding of AR function in PCa cell lines and tissues (Pomerantz et al., 2015; Sharma et al., 2013; Stelloo et al., 2018), much less is known of the mechanisms underlying AR-mediated transcriptional regulation in normal prostate cells. Thus, to define the epigenomic response to AR activation, we generated 3D organoid models of murine prostate epithelial cells (Karthaus et al., 2014). Upon establishment, these prostate organoids faithfully mimicked prostate gland architecture (Karthaus et al., 2014) and immunohistochemical features (Fig. S1a). To catalogue the function of AR in the normal prostate, we assessed genome-wide changes in transcriptional profiles (RNA-seq), chromatin accessibility by assay for transposase-accessible chromatin (ATAC-seq), and cistromes of H3K4me2, AR, and FOXA1 by chromatin immunoprecipitation sequencing (ChIP-seq) post-treatment with 10 nM DHT (Fig. 1a).

**Fig. 1.**
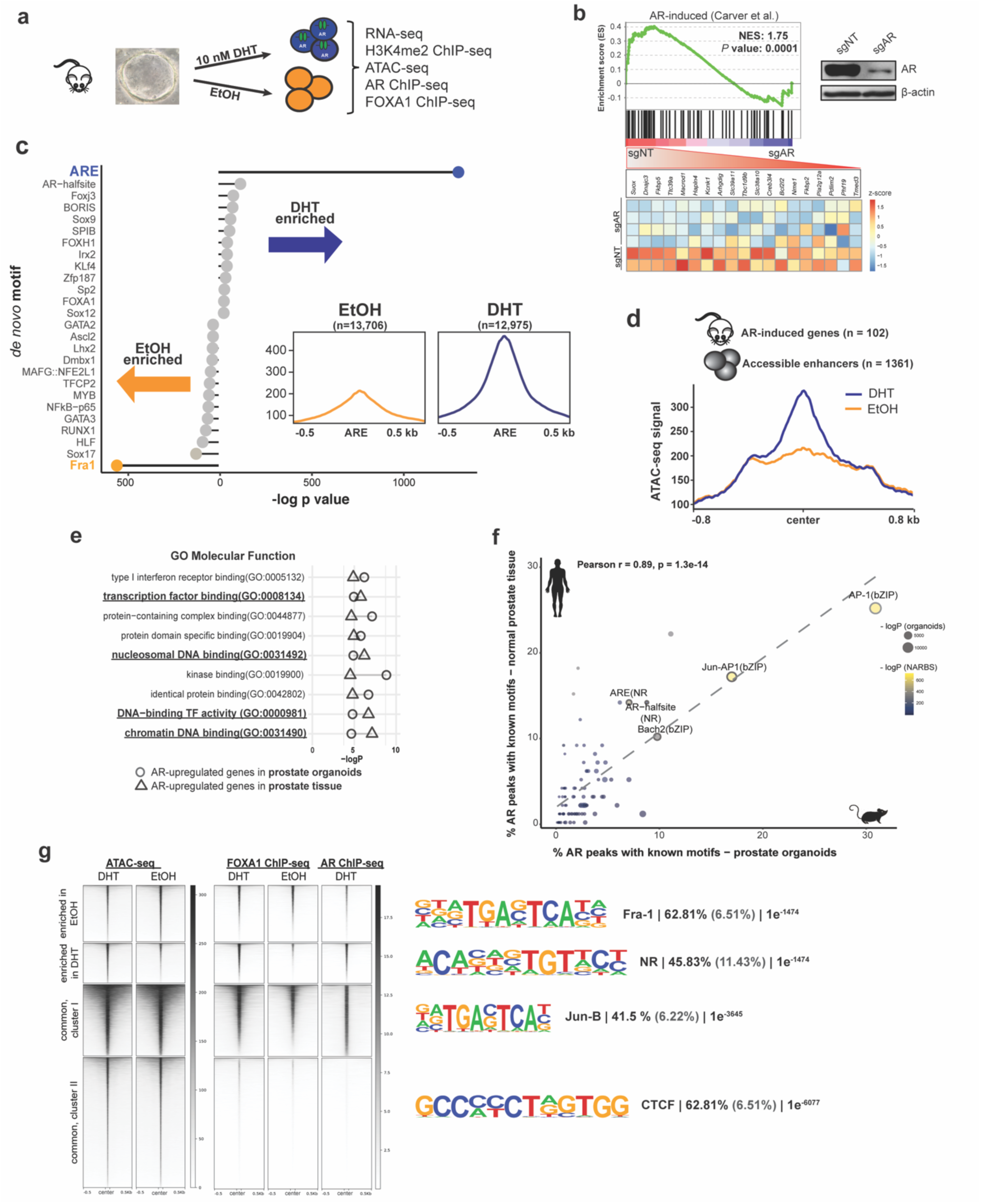
DHT-induced changes in chromatin accessibility correspond to AR-transcriptional response. **(a)** Schematic representation of treatments and obtained datasets. **(b)** GSEA and leading edge analysis of AR activity in normal prostate organoids. Upper right corner: Immunoblot of murine organoid CRISPR-Cas9 cells with non-targeting (NT) or *Ar*-specific sgRNA. **(c)** Lolipop plot representing known motif enrichment in vehicle-treated and androgen-treated murine organoids. **(d)** Histogram depicting accessibility signal from EtOH-treated and DHT-treated organoids over the center of regulatory elements within 1 Mb of AR-induced genes from prostate tissue. **(e)** Cleveland plot indicating results of gene ontology (GO) analysis of AR-induced genes in prostate organoids (circle) and prostate tissue (triangles) integrated with accessible regions identified after androgen stimulation of organoid cells. **(f)** Known motif analysis of AR ChIP-seq datasets from normal mouse organoids (x-axis) and human normal prostate tissue (n = 7; GEO: GSE70079; y-axis) demonstrating enrichment for similar motifs (r = 0.89, p < 0.001). The dotted line represents a regression line. Circle size represents the -log P-value of motif enrichment in AR ChIP-seq peaks in normal murine prostate organoids and circle color the motif enrichment significance in AR binding sites in normal prostate tissue (NARBS). **(g)** ATAC-seq, FOXA1, and AR ChiP-seq signals over the regions that were identified to be more open before (enriched in EtOH) or after androgens (enriched in DHT). The tornado plots also cover the regions whose accessibility is not DHT-dependent (common). The most enriched *de novo* motifs per cluster are depicted on the right, along with the percentage of target (black) and background (grey) regions and the resulting p-value.

Gene expression data (Fig. S1b) showed upregulation of known murine AR-regulated genes (Carver et al., 2011) by androgens in intact organoids, but not those with CRISPR-mediated deletion of Ar (sgAr). We successfully validated the transcriptional response by performing GSEA on AR-regulated genes in prostate tissue (Carver et al., 2011) (Fig. 1b), confirming a relevant transcriptional response in our organoid model system (Fig. S1 b-f).

To define AR-driven alterations in the chromatin landscape in normal prostate epithelial cells, we performed ATAC-seq, as a proxy for genetic regulatory element activity (Fig. S1g-h and Tables S1 and S2). We identified 26,681 high-confidence differentially accessible peaks between DHT-treated and vehicle-treated prostate organoids (FDR < 0.01, Fig. S1i). Next, we performed *de novo* motif analysis of the DHT-enriched versus depleted accessible elements, revealing significant enrichment of the androgen response element (ARE) motif in DHT-treated cells, which was associated with an increase in accessibility at these sites post DHT treatment (Fig. 1c). To connect the DHT-responsive regulatory elements to AR-target genes in prostate tissue, we profiled the ATAC-seq signal of androgen-regulated genes in the mouse prostate (Carver et al., 2011) (Fig. 1d). Subsequently, we performed functional enrichment analysis of the ATAC-seq defined DNA regulatory elements and associated AR-induced genes in both prostate organoids and tissue, with “transcription factor binding” and “chromatin (nucleosomal) binding” among the enriched categories (Fig. 1e). Overall, our ATAC-seq analysis confirmed a robust and biologically relevant response to androgens at the chromatin level.

Next, to nominate the accessible peaks directly regulated by AR binding, we profiled AR localization genome-wide after DHT stimulation. Genome-wide localization of AR by ChIP-seq resulted in 42,206 AR peaks (Table S1). Importantly, our motif analysis found that the canonical ARE was the most enriched motif in this dataset, confirming the specificity of our signal. Further analysis comparing motif enrichment against that of AR in human prostate tissue (Pomerantz et al., 2015) uncovered a highly similar motif enrichment pattern (r = 0.89, p < 0.001) (Fig. 1f), suggesting our models faithfully recapitulate critical features of AR signaling.

Integrative analysis across data types revealed distinct clusters of regulatory sites either unique to baseline or AR stimulated conditions, or common between these, with associated dominant transcription factors motifs (Fig. 1g, S1e-j). While nuclear hormone receptor motifs were most enriched in dynamic peaks associated with DHT stimulation (cluster 2), regions with chromatin accessibility enriched in basal conditions (cluster 1) or common (cluster 3) were associated with AP-1 motifs (Fra-1 and Jun-B). AP-1 motifs are also common in AR ChIP-seq from normal human prostate tissue (Figure 1f), consistent with an AR-directed role in non-transformed prostate cells.

Together, these data demonstrate that androgen stimulation of prostate organoids results in chromatin and transcriptional changes concordant between both murine and human prostate tissues and provides needed insight into the molecular functions of AR in a genetically normal context in addition to being a valuable resource to study early events which contribute to AR driven prostate tumorigenesis. In addition, it is a valuable resource to study early events which contribute to AR driven prostate tumorigenesis.

### SPOP mutation alone alters the landscape of accessible enhancers in response to androgen

*SPOP* mutations are present in about 10% of PCa (Barbieri et al., 2012; Cancer Genome Atlas Research, 2015; Li et al., 2020), occur early in the natural history of the disease, are relatively prostate-specific, and affect AR activity through deregulation of multiple substrates (Blattner et al., 2017; Geng et al., 2013; Theurillat et al., 2014). We previously developed a transgenic mouse with Cre-dependent, conditional expression of SPOP-F133V (Fig. 2a) (Blattner et al., 2017). In prostate organoids infected with the Cre virus and vector, we examined the impact of an *SPOP* mutation on the epigenomic and transcriptional landscapes after androgen stimulation (Fig. 2b).

**Fig. 2.**
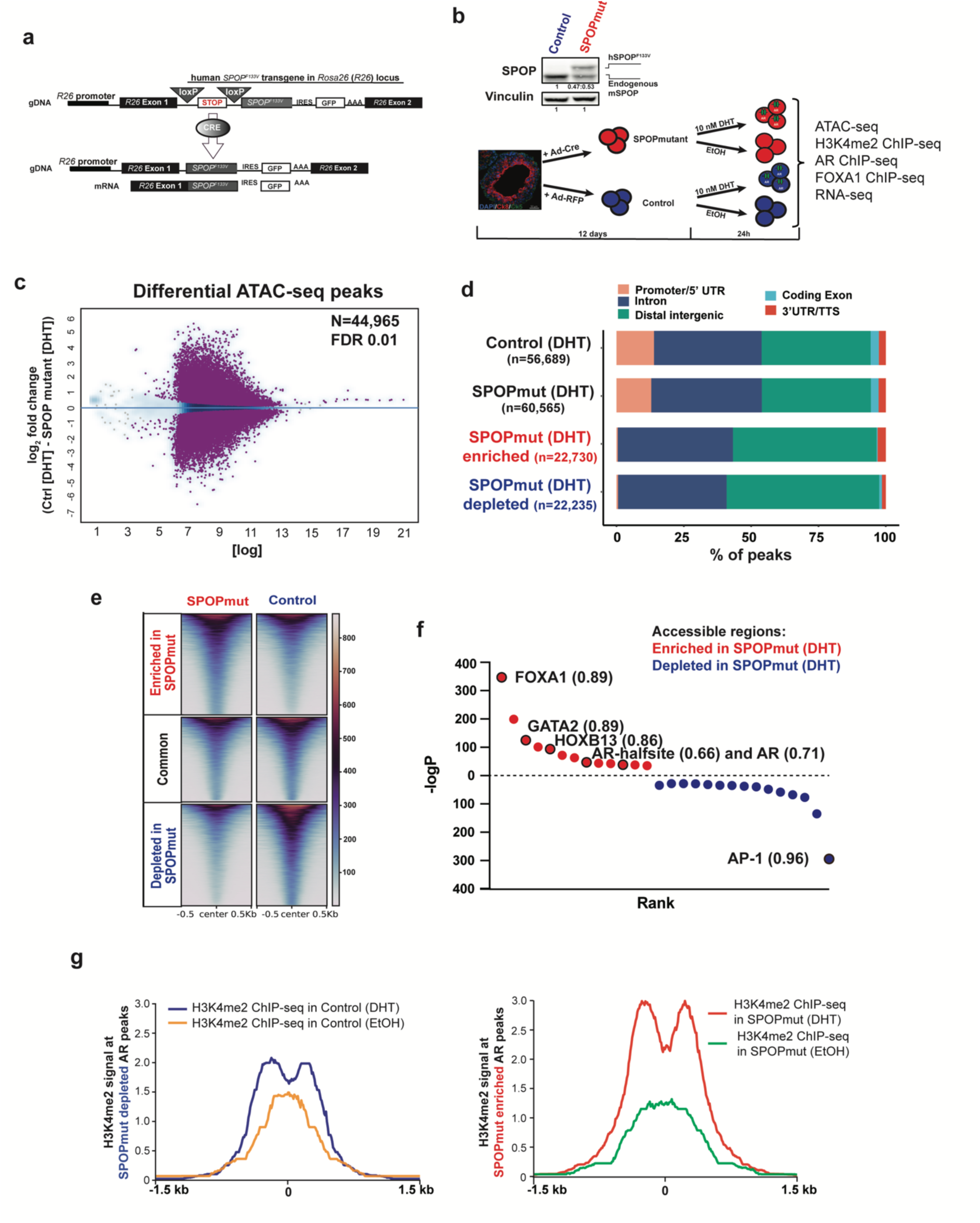
Extensive remodelling of enhancers in SPOP-mut organoids. **(a)** Schematic of conditional *SPOP-F133V* construct in the *Rosa26* (*R26*) locus (top) and of the expressed transgenic transcript after Cre introduction (bottom). **(b)** Illustration of the generation of the organoid lines and treatments in addition to all the performed experiments. Up: Immunoblot showing physiological levels of human SPOP-F133V (top band) in comparison to endogenous mouse SPOP protein (bottom band). **(c)** Differential accessible peaks (purple) between Control (n=4) and SPOP-mut (n=4) cells at FDR 0.01. **(d)** Genomic annotation of consensus and differential peaks showing predominant enhancer location of *SPOP* mutant enriched and depleted accessible elements. **(e)** Heatmap of accessible enhancers showing differential accessibility in SPOPmut enriched and depleted peaks. **(f)** *De novo* motif analysis of SPOPmut enriched and depleted accessible DNA regulatory regions. Enriched motifs are plotted according to their rank and *P* value. Values next to the motif are the best match motif score (max. is 1). **(g)** H3K4me2 signal at the enhancers from Control and *SPOP* mutant organoids with and without DHT stimulation at *SPOP* mutant depleted AR peaks (left), and *SPOP* mutant enriched AR peaks (right).

We profiled the chromatin accessibility landscape of androgen-stimulated prostate organoids expressing mutant (Fig. S2a-c) and wildtype SPOP using ATAC-seq (Fig. S1g-i). We found > 44.9k differentially accessible sites (Fig. 2c), the majority of which were not promoter-associated, but at distal intergenic and intronic regions, consistent with enhancers (Fig. 2d). Dynamic accessible peaks enriched in *SPOP* mutant organoids (Fig. 2e) were enriched for motifs of TFs important for AR-driven prostate tumorigenesis (Fig. 2f), including FOXA1 and HOXB13 (Pomerantz et al., 2015). These data support that the chromatin landscape of *SPOP*-mutant prostate cells is remodelled (Fig. S2d-e), especially at androgen-responsive enhancer regions associated with lineage-specific, oncogenic transcription factors.

H3K4me2-marked nucleosomes go through remodelling upon AR binding to chromatin (He et al., 2010). Upon androgen stimulation, the H3K4me2 nucleosome at the center of ARE is evicted, and subsequently, AR-binding sites are flanked with H3K4me2 nucleosomes. The same bimodal distribution before and after androgen stimulus was shown in the case of AR-pioneering factor FOXA1 as well (He et al., 2010). Consistent with our chromatin accessibility data, there was more pronounced chromatin remodelling (measured by H3K4me2) at AR bound enhancers in *SPOP*-mutant organoids (Fig. 2g) that also showed a higher frequency of FOXA1 and joint FOXA1:AR motifs.

### SPOP mutation drives an oncogenic AR-cistrome in prostate organoids

Oncogenic reprogramming of the AR cistrome is a fundamental feature of prostate cancer (Augello et al., 2019; Pomerantz et al., 2015; Sharma et al., 2013; Stelloo et al., 2018), but which specific alterations can drive this phenotype, and what stage of the disease it occurs, remains unclear. We sought to test the hypothesis that a single early event like *SPOP* mutation could impact reprogramming of the AR cistrome to mimic PCa. AR ChIP-seq (Tables S1 and S2) with a validated AR antibody, revealed 13,598 differential AR binding sites (FDR < 0.01) between control and *SPOP-*mutant organoid lines (Fig. 3a). In *SPOP-*mutant organoids, the AR cistrome was enriched for the motifs of well-established regulators of AR function associated with tumorigenesis, including FOXA1 and HOXB13 (Fig. 3b). Strikingly, the motif enrichment pattern in *SPOP-*mutant organoids compared to controls was remarkably similar to that of previously reported in AR cistromes of human prostate adenocarcinoma (PRAD) samples (Pomerantz et al., 2015) as compared to normal prostate tissue (Fig. S3a-b). Consistent with ATAC-seq results, AR peaks in *SPOP-*mutant prostate organoids harbored a higher density of canonical AR and FOXA1 motifs (Fig. 3c-d). To validate the relevance of this shift in the TF motif occurrence, we analyzed the AR cistromes of human primary prostate carcinoma tissue samples (Stelloo et al., 2018) (Fig. 3e). AR peaks from human *SPOP* mutant cancers had higher FOXA1 motif density (Fig. 3f), associated with a shift in the binding of FOXA1 towards the center of AR peaks (Fig. 3g), consistent with our murine organoid model. Collectively, these data suggest that that *SPOP* mutation alone alters the androgen-responsive enhancer landscape, drives an AR cistrome consistent with PCa, and nominates FOXA1 as a potential mediator of this phenotype.

**Fig. 3.**
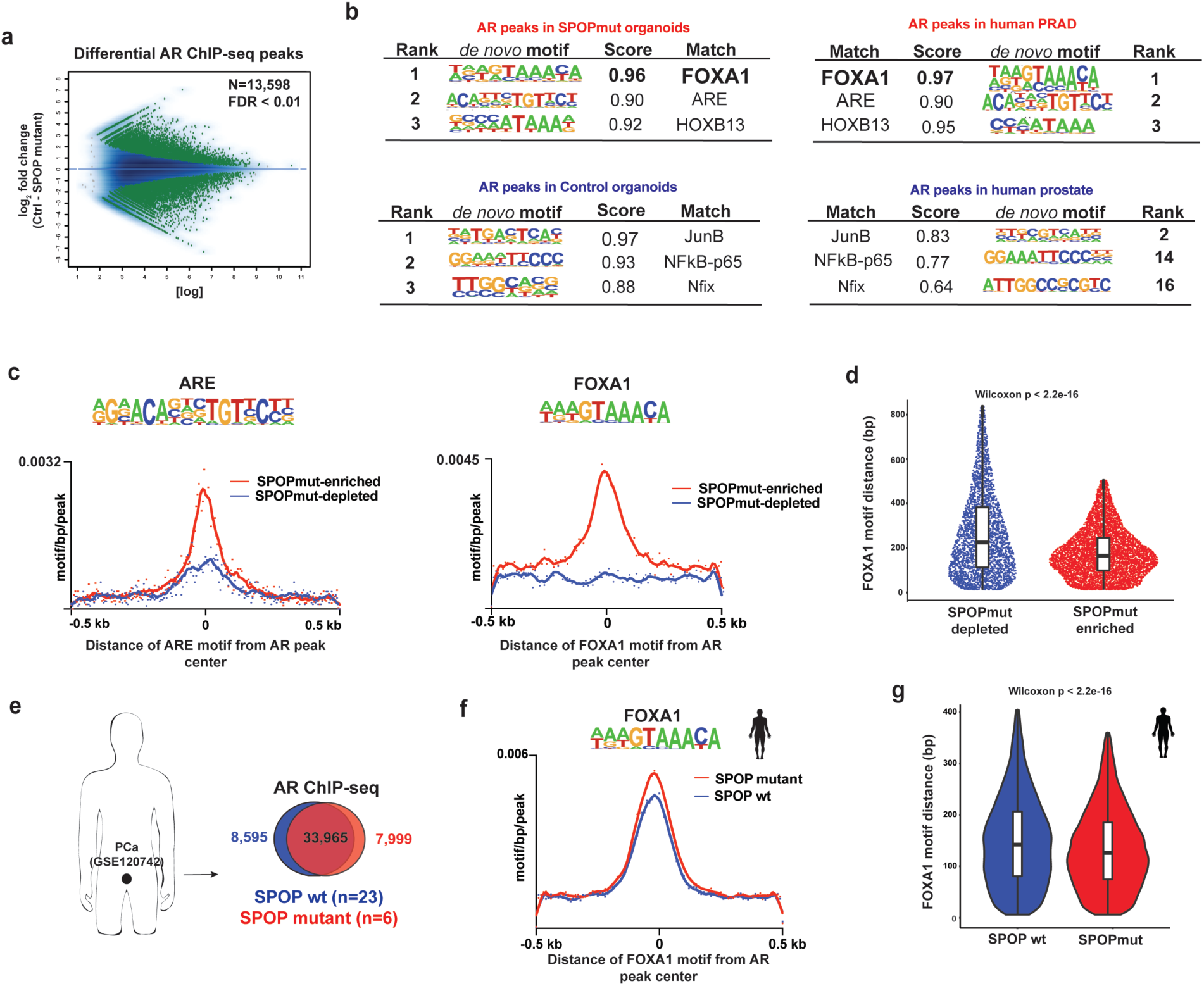
AR cistrome from *SPOP*-mutated cells is enriched for FOXA1 motifs. **(a)** MA plot showing AR differential sites (green dots) between Control and SPOP-F133V expressing organoids using two biological replicates per condition. **(b)** *De novo* motif analysis of AR cistrome in SPOPmut and Control organoids (right). Left: the same type of motif analysis performed on human PRAD (Pomerantz et al., 2015). **(c)** Histograms of ARE (left) and FOXA1 (right) motif densities within a 500 bp window around the center of each AR peak. Input files were from *SPOP* mutant enriched (red line) and depleted (blue depleted) AR ChIP-seq peaks. **(d)** Sina plot – distances of FOXA1 motif from the center of each AR peak. Input files were as in (c). **(e)** AR ChIP-seq data from human prostate cancer tissues (Stelloo et al., 2018) (GSE120742). Peaks coming from either PRAD with *SPOP* mutation or wt *SPOP* were merged and intersected. **(f)** Histogram of FOXA1 motif density in AR binding sites from human prostate cells with SPOP-F133V (red line) or wt SPOP (blue). **(g)** The distance of the FOXA1 motif from the center of AR ChIP-seq peaks in each prostate cancer group.

### Increased FOXA1 binding in SPOP mutant tumors

FOXA1 acts as a pioneer factor, binding and opening condensed chromatin via its winged-helix DNA binding domains (Cirillo et al., 2002), facilitating AR-mediated transcriptional control (Gao et al., 2003). To better define the role of FOXA1 nominated by motif enrichment in a subset of SPOPmut-specific AR peaks (Fig. 3), we assessed genome-wide FOXA1 binding. *SPOP* mutant organoids and controls showed 27,168 differential FOXA1 peaks (Fig. 4a, Table S1). Moreover, the highest FOXA1, FOXA1:ARE, and ARE motif densities occurred within the SPOPmut-specific FOXA1 cistrome (Fig. 4b). Next, we interrogated AR binding at FOXA1-bound enhancers and found profound enhancement of AR signal at FOXA1-bound accessible enhancers specific for *SPOP* mutant prostate cells (Fig. 4c-d). These data implicate a role for FOXA1 downstream of *SPOP* mutation in driving pro-tumorigenic changes at a subset of AR binding loci.

**Fig. 4.**
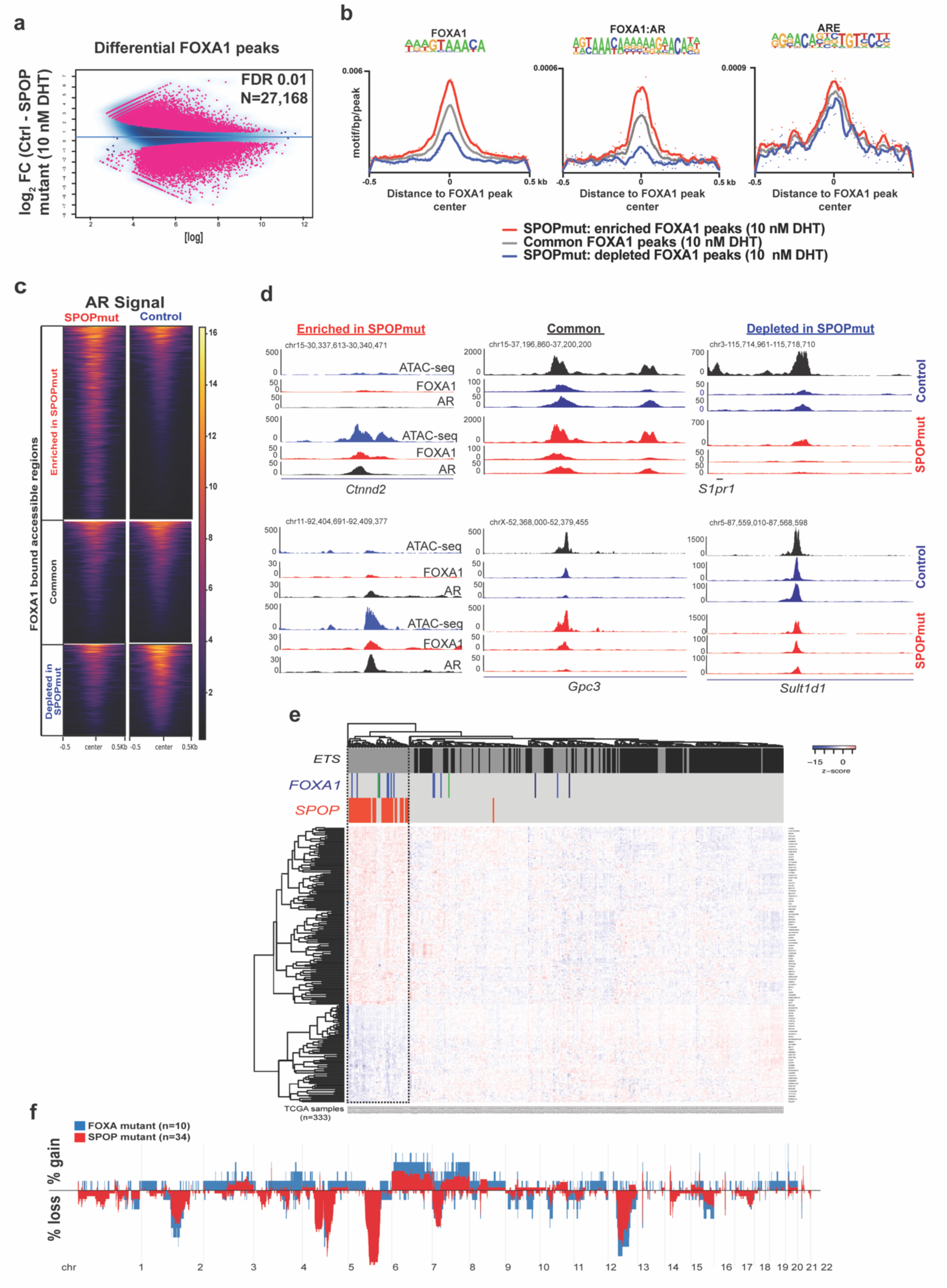
Increased binding of FOXA1 at *SPOP* mutant-specific AR sites. **(a)** Differential FOXA1 genome-wide binding in Control (n=2) and *SPOP* mutant (n=2) organoids upon AR activation. **(b)** Motif densities within the center of FOXA1 peaks enriched (red) or depleted (blue) in *SPOP* mutant murine organoids. Gray - The signal at regions where FOXA1 binding intensity does not change. **(c)** AR binding signal at FOXA1-bound DNA regulatory elements that are common between the lines or *SPOP* mutant enriched or depleted. (**d**) Representative Genome Browser snapshots of regions analysed in (c). **(e, f)** Relationship between *SPOP*-mutant and *FOXA1*-mutant subclasses of PRAD. **(e)** Classification of PRAD using the *SPOP*-mutant transcriptional classifier (Liu et al., 2018). *FOXA1* mutations are depicted in blue (FKHD – DNA binding domain), green (N-terminal domain) and black (truncations after FKHD domain). **(f)** Gene copy number profiles of these molecular subclasses of PCa.

We next looked in human prostate cancers for evidence to support a relationship between *SPOP* mutation and FOXA1. *FOXA1* is recurrently mutated in localized prostate cancer, with forkhead (FKHD) domain mutations altering pioneering activity being the most common (Adams et al., 2019; Barbieri et al., 2012; Cancer Genome Atlas Research, 2015; Parolia et al., 2019). We hypothesized that if *SPOP* mutations acted in part through modulation of FOXA1 activity, then *SPOP*-mutant and *FOXA1*-mutant tumors would display similar features. When applying a previously reported transcriptional classifier for *SPOP* mutation (Liu et al., 2018), the majority of *FOXA*1 mutant tumors cluster together with *SPOP* mutant tumors (Figure 4e). In addition, tumors harbouring *SPOP* mutations and those with *FOXA1* alteration display similar somatic copy number alterations (Figure 4f), suggesting common collaborating events.

### Gene expression alterations in SPOP mutant prostate organoids

To define whether the changes in accessibility and FOXA1 and AR binding are reflected in transcriptional response to androgens in *SPOP* mutant cells, we performed RNA-seq (Fig. S4a-b). Importantly, AR induced and repressed distinct genes in Control and SPOPmut cells (Fig. 5a).

**Fig. 5.**
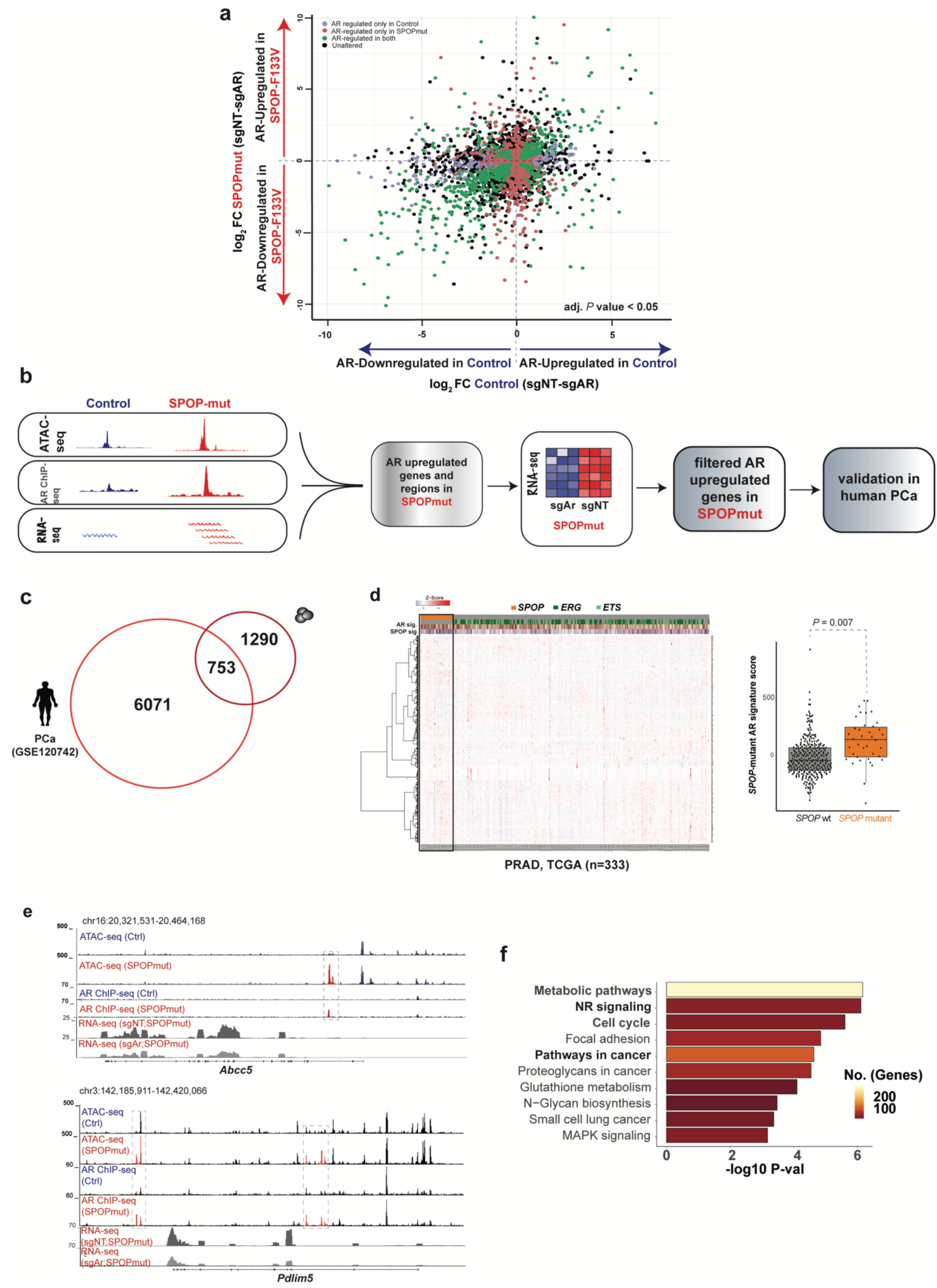
Nominating novel AR-regulated genes in the context of SPOP mutation. **(a)** Plot showing differentially expressed genes (adj. *P* value < 0.05) regulated by AR in Control cells (x-axis; blue) and *SPOP*-mut cells (y-axis, red). **(b)** GSEA hallmark enrichment of AR-responsive genes in Control (blue) and *SPOP* mutant (red) cells with plotted GSEA NES values. **(c)** Computational steps to identify AR-upregulated genes specific to *SPOP* mutant human and murine cells. **(d)** Overlapping AR-regulated genes in *SPOP*-mutant human PCa (Stelloo et al., 2018) and murine cells. **(d)** Heatmap - Gene expression (n=753) in human PRAD samples (Cancer Genome Atlas Research, 2015). Boxplots – PCa samples with higher expression of nominated genes (n=753) have higher *SPOP*-mutant signature score. **(e)** Representative Genome Browser tracks of AR-regulated regulatory regions and transcripts in *SPOP*-mut organoids. **(f)** Functional enrichment of androgen-induced genes in murine *SPOP* mutant cells.

We next sought to leverage multiple levels of data to define genes that are AR-regulated specifically in the context of *SPOP* mutation (Fig. 5b). Integrative analysis of ATAC-seq, AR ChIP-seq, and RNA-seq data nominated genes linked to dynamic regulatory events in *SPOP* mutant mouse organoids, and dependent on AR for induction (n=2043). Of these, 753 genes were also upregulated in *SPOP* mutant human prostate cancers (Fig. 5c), and these were used to generate an *SPOP*-mutant AR signature score (Fig. 5d). By correlating across datatypes, we identified AR-driven, SPOP mutation specific transcriptional regulatory events, with increased AR binding, increased accessibility, and AR-dependent increased gene expression in genes such as *Abcc5* and *Pdlim5*, which has been previously linked to prostate cancer pathogenesis (Liu et al., 2017). Functional categorization of these *SPOP*-mutant specific AR-regulated genes revealed enrichment for metabolic pathways, Nuclear receptor signaling, and cell cycle, consistent with an AR-dependent role in prostate tumorigenesis (Fig. 5f). Together, we conclude that the SPOPmut associated changes in accessibility and transcription factor binding drive a distinct AR-dependent gene expression program, one more consistent with tumorigenesis.

### Critical dependency of SPOP mutant prostate epithelium on AR signaling

Having observed that *SPOP* mutation alters the androgen-responsive chromatin landscape and transcriptional program, we next examined whether *SPOP* mutant prostate organoids showed increased dependency on AR signaling. *SPOP* mutant organoids showed increased sensitivity to AR inhibition with bicalutamide compared to controls (Fig. 6a), with the proliferation of the SPOPmut significantly more sensitive to anti-androgen treatment both in 2D and 3D growth conditions (Fig. S5a-b). Given the importance of FOXA1 in the epigenomic alterations in *SPOP*-mutant cells, we hypothesized that FOXA1 was necessary for the AR dependency driven by *SPOP* mutation as well. Transient downregulation of FOXA1 further increased the sensitivity of SPOPmut organoids to AR inhibition (Fig. 6b).

**Fig. 6.**
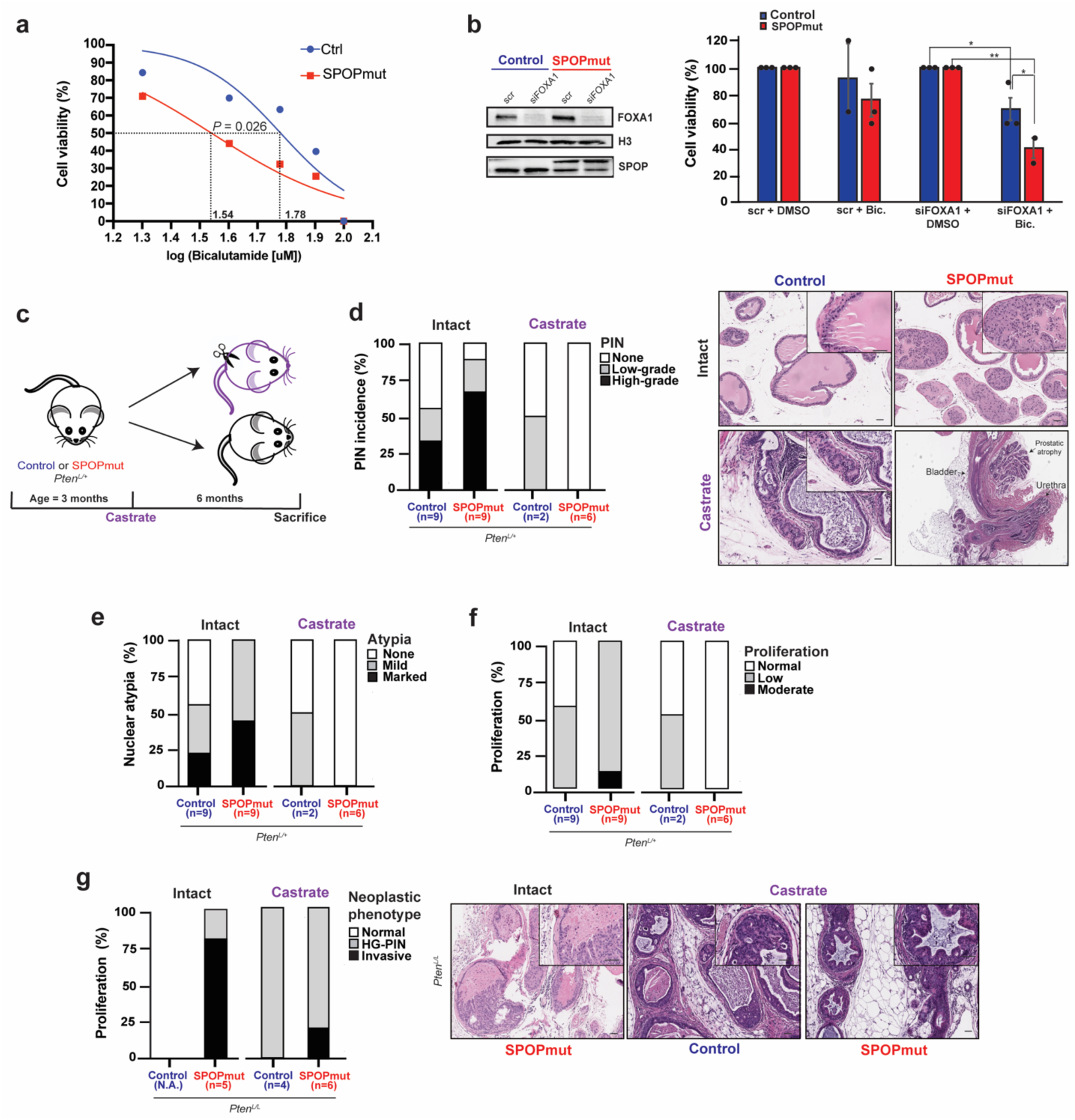
The distinct phenotype of *SPOP* mutations shows dependency on functional AR signaling. **(a)** The dose-response curve (log scale) of Control and SPOPmut murine prostate organoids in response to bicalutamide. **(b)** Immunoblot of control and SPOPmut murine prostate cells after siRNA-mediated downregulation of FOXA1 (top) and cell viability of these cells after 20 μM bicalutamide treatment (bottom). **(c)** Schematic of castration experiment. **(d-f)** Incidence of PIN, nuclear atypia and degree of proliferation in Control and SPOPmut mice before and after castration. Representative H&E images are shown on the right and in Fig S5c-d. Scale bar, 50 mm. **(g)** Incidence of neoplastic phenotypes (HG-PIN and invasive features such as necrosis, stromal response and/or frank invasion) in Control and *SPOP-F133V* mice with representative images shown on the right. Scale bar, 50 mm.

To investigate the role of AR signaling in a mouse model of *SPOP-F133V*, we performed a pathologic analysis of castrate and hormonally intact mice in a background of heterozygous *Pten* loss (*PbCre^+^;Pten^L/+^;Rosa26^F133V^*). We have previously shown that *SPOP* mutation alone is insufficient to drive tumorigenesis in mouse prostate (Blattner et al., 2017). However, in a conditional *Pten* deleted background, *SPOP* mutation results in more aggressive neoplastic histology, including high-grade PIN (HG-PIN) in *Pten^L/+^* mouse prostates and invasive carcinoma in *Pten^L/L^*. We castrated *SPOP* mutant (*PbCre^+^;Pten^L/+^;Rosa26^F133V^*) and control (*PbCre^+^;Pten^L/+^*) mice at 8 weeks, sacrificed them at 9 months and performed a histological examination of their prostates compared to age-matched uncastrated controls (Fig. 6c). Castration reversed the increased amount of HG-PIN, nuclear atypia, and proliferation seen in *SPOP* mutant prostates compared to controls (Fig. 6d-f and S5c-d). In the setting of homozygous *Pten* loss, *SPOP-*mutant mice (*PbCre^+^;Pten^L/L^;Rosa26^F133V^*) develop invasive, poorly differentiated carcinomas, while control mice (*PbCre^+^;Pten^L/L^*) developed only diffuse HG-PIN (Blattner et al., 2017). In this background, we found that *SPOP* mutant castrated mice develop features of invasive disease less frequently than hormonally intact mice, while the diffuse, proliferative HG-PIN in both *SPOP* mutant mice and *Pten^L/L^* controls (Fig. 6g) remains relatively castration-resistant, consistent with prior reports (Blattner et al., 2017; Mulholland et al., 2011). Together, these data show that castration results in the reversal of *SPOP* mutant phenotypes *in vivo*, suggesting that these phenotypes are driven by AR signaling.

### SPOP mutation is associated with improved response to AR-targeted therapies

Given the prominent role of AR signaling in *SPOP* mutant preclinical models, we investigated the impact of *SPOP* mutation on patient response to therapies targeting AR in multiple clinical scenarios. Using a previously developed transcriptional classifier for the *SPOP* mutant subclass (Liu et al., 2018), we examined data derived from the Decipher Genomics Resource Information Database (GRID) registry (ClinicalTrials.gov identifier: NCT02609269) (Liu et al., 2018). In a pooled retrospective cohort of 1,626 patients *SPOP* mutant tumors were associated with improved metastasis (MET)-free survival in patients who received androgen deprivation therapy (ADT) after radical prostatectomy, but no difference without ADT (Fig. 7a and Table S3). Next, we hypothesized that preferential sensitivity to ADT would manifest with relative depletion of *SPOP* mutant tumors in castrate-resistant prostate cancer compared to hormone-sensitive disease. Comparing 333 primary prostate adenocarcinomas (Cancer Genome Atlas Research, 2015) and 414 metastatic CRPC samples (Abida et al., 2019), *SPOP* mutations were significantly lower in CRPC (Fig. 7b), suggesting a response to ADT. To rule out an effect of primary vs. metastatic disease in this analysis, we compared metastatic hormone-sensitive PCa (mHSPC) to metastatic hormone-resistant PCa (mCRPC) within the same cohort (Abida et al., 2017), and again observed significant underrepresentation of *SPOP* mutation in mCRPC, consistent with the sensitivity of *SPOP* mutant metastatic prostate cancers to ADT (Fig. 7c). Finally, we examined the response of patients with mCRPC patients to AR-targeted therapy (abiraterone and enzalutamide) (Abida et al., 2019). *SPOP* mutation was associated with a longer time on treatment in these patients, suggesting that AR-targeted therapies may be a more durable treatment option in *SPOP* mutant prostate cancer than in other subtypes (Fig. 7d). Overall, all these results together show a favourable outcome of *SPOP* mutant PCa to AR-targeted therapy across multiple clinical scenarios and suggest that *SPOP* mutation may be a predictive biomarker for these therapeutic regimens.

**Fig. 7.**
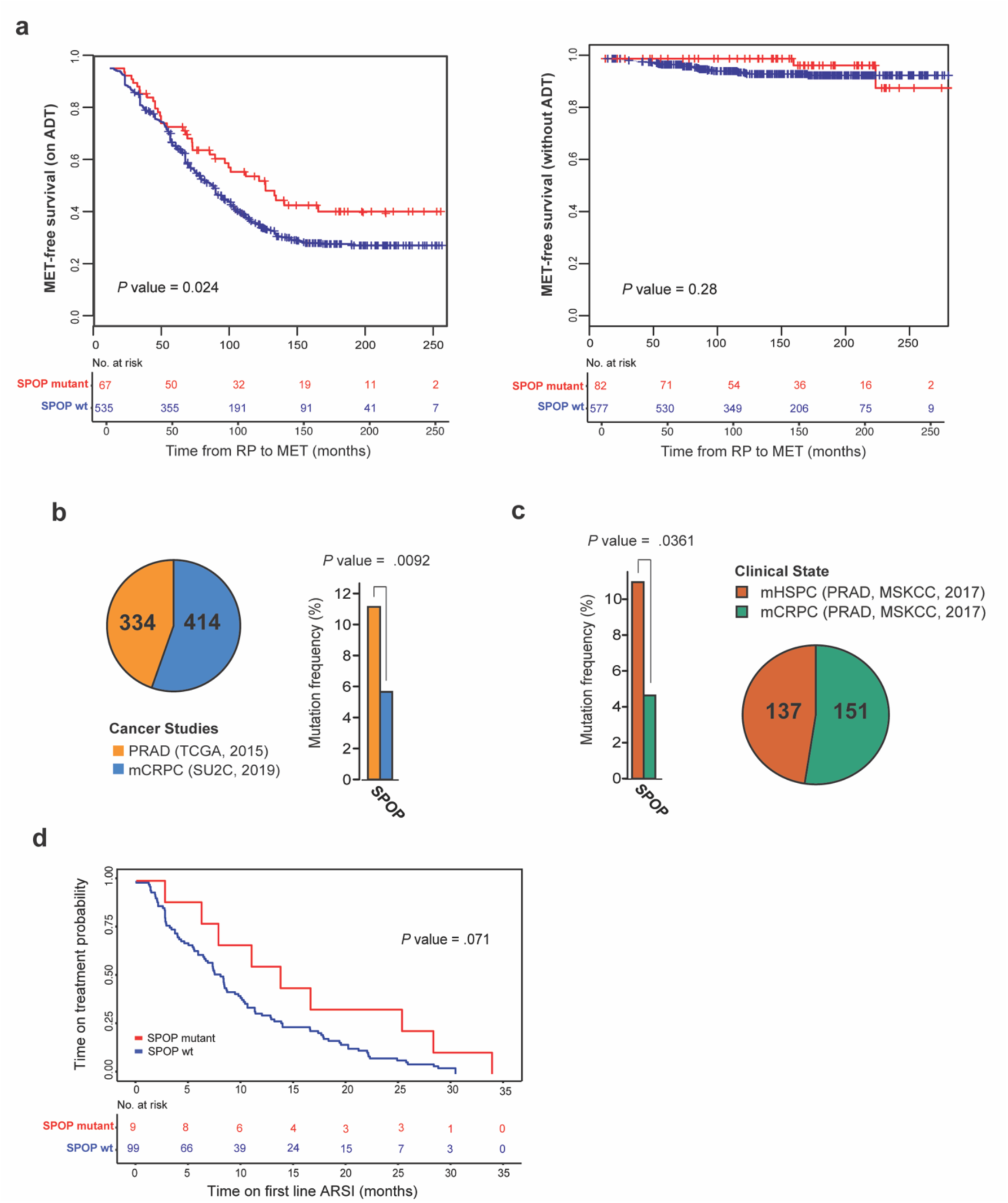
Sensitivity of *SPOP* mutant PCa to therapies targeting AR signaling. **(a)** Significant clinical outcome difference between *SPOP* mutant and wild-type PCa via Kaplan-Meier analysis of metastasis (MET)-free survival for patients on androgen deprivation therapy (ADT) (left) or without receiving ADT (right). **(b)** The occurrence of *SPOP* mutation in primary PCa (Cancer Genome Atlas Research, 2015) and metastatic (mCRPC) samples (Abida et al., 2019). **(c)** When metastatic prostate tumors (Abida et al., 2017) are compared – the hormone-sensitive ones (mHSPC) have a higher frequency (p < 0.0361) of *SPOP* mutation than the resistant (mCRPC) ones. **(d**) Kaplan-Meier analysis of *SPOP* wild-type and *SPOP*-mutated mCRPC tumors (27) showing time on treatment with first-line androgen signaling inhibitors (ARSI).

## Discussion

The androgen receptor (AR) is the central determinant of prostate tissue identity and differentiation (Cunha et al., 2004), controlling normal, growth-suppressive prostate-specific gene expression (Schiewer et al., 2012). However, it is also a central driver of prostate tumorigenesis, becoming “hijacked” to drive oncogenic transcription (Schiewer et al., 2012). Importantly, it also remains the key therapeutic target for PCa, even in advanced castrate-resistant prostate cancer (CRPC) (Watson et al., 2015). However, how AR function is altered to convert it to an oncogenic factor, and at what steps in the process of tumorigenesis this occurs, remain poorly defined. Here, we show that a single early event in the natural history of prostate cancer is sufficient to drive oncogenic alterations in the AR-directed chromatin landscape and transcriptional program.

It has been long recognized that AR is required for the development and function of the normal prostate (Heinlein and Chang, 2004). However, how AR reshapes the landscape of chromatin to execute its transcriptional program, particularly in genetically normal cells, remains unclear. Here, using genetically normal, physiologically relevant prostate organoids, we defined the genome-wide effect of androgens on chromatin accessibility, H3K4me2, AR, and FOXA1 binding and subsequent changes in the transcriptome. These data provide insight to transcriptional dynamics of normal AR function, and demonstrate that these genetically pliable models can recapitulate the epigenomic and transcriptomic changes of mouse and human tissue, credentialing a valuable tool to investigate AR activity.

Missense mutations in *SPOP* are the most common mutations in primary prostate adenocarcinoma, occur early in the natural history of the disease and affect multiple substrate proteins associated with AR-mediated transcriptional regulation (Blattner et al., 2017; Bouchard et al., 2018; Geng et al., 2013; Geng et al., 2014; Theurillat et al., 2014). Here, we show that *SPOP* mutation alone, in otherwise normal prostate organoids, is sufficient to induce dramatic changes in accessibility of regulatory elements. Furthermore, *SPOP* mutation was sufficient to reprogram the AR cistrome to one with striking resemblance to human PCa. Interestingly one other study reported different effects on the AR cistrome in response to *SPOP* mutation introduced into established cell lines with multiple alterations (Copeland et al., 2019), with expansion of the AR cistrome towards both normal and tumorigenic sites. The more profound shift we report here may reflect the utility of genetically normal models in studying initiation events, and suggest that oncogenic reprogramming of the AR transcriptional program may be a very early event in the natural history of PCa, at least in selected subtypes.

FOXA1 is a key pioneering factor for AR and HOXB13 transcription factors, facilitating binding to at PCa-specific AR bound-sites. Furthermore, FOXA1 and HOXB13 overexpression transforms AR cistrome from normal, immortalized cell lines into an oncogenic one (Copeland et al., 2019; Pomerantz et al., 2015). Our study points to a significant role for FOXA1 during the reprogramming of AR cistrome in *SPOP* mutant prostate organoids, consistent with observations in human tumors.

The data here suggest that oncogenic reprogramming of AR-driven transcription is the major mechanism underlying the oncogenic function of *SPOP* mutation in prostate cancer. Consistent with this, we also find that functionally, *SPOP* mutant model systems show high dependency on this pathway, and in clinical data, *SPOP* mutant human prostate cancers show increased sensitivity to androgen-targeted therapy. Strikingly, our analysis of the data from multiple clinical scenarios supports this hypothesis: patients receiving adjuvant or salvage ADT after prostatectomy (Fig. 7a) and second generations AR inhibitors with CRPC (Fig. 7d), as well depletion of *SPOP* mutation in CRPC compared to both primary and hormone sensitive metastatic disease (Fig. 7b-c), consistent with reports of favourable outcomes for *SPOP* mutant cancers with therapies targeting AR in both CRPC (Boysen et al., 2018) and hormone sensitive metastatic disease (Swami et al., 2020). However, credentialing *SPOP* mutation as a clinically actionable predictive biomarker for AR-directed therapies will require clinical trials designed for this purpose, likely in each specific clinical scenario. Consistent with this, Tewari and colleagues report that *SPOP* mutation is a truncal alteration in human prostate cancers that show exceptionally robust response to neoadjuvant androgen-targeted therapy (in review). Together, these clinical data suggest that the AR-dependent effects of *SPOP* mutation are not unique to a single clinical scenario, but may extend to multiple stages of prostate cancer, broadening the potential impact.

## Methods

### Transgenic mouse model

Weill Cornell Medicine (WCM) Institutional Care and Use Committee approved all the mouse studies under protocol 2015-0022. Transgenic mice had prostate-specific expression of human *SPOP-F133V* in *Rosa26* locus (*PbCre;R26^F133V/WT^* or *PbCre;R26^F133V/F133V^*) with or without *Pten* deletion (*PbCre;R26^F133V/WT^*;*Pten^+/+^*, *PbCre;R26^F133V/WT^*;*Pten^L/+^, PbCre;R26^F133V/F133V^*;*Pten^L/+^* or *PbCre;R26^F133V/WT^*;*Pten^L/L^*)(Blattner et al., 2017). TransNetYX performed mouse genotyping by using published primers (Augello et al., 2019; Blattner et al., 2017).

### Immunohistochemistry

Whole murine prostates were processed as previously described(Augello et al., 2019). Staining was done at The Translational Research Program (TRP) of the Department of Pathology and Laboratory Medicine at WCM. Ki67 (ab16667) was used for staining.

### Murine Organoid Line Generation and CRISPR experiments

Prostate from the *PbCre4* negative; *Rosa26^F133V^* mice were harvested at 1-2 months of age and processed and grown as 3D Matrigel culture as previously described (Drost et al., 2016) with the exception that the organoid media contained 5 ng/mL EGF. Lines were generated by infection with Adeno-RFP (Control cells) or Adeno-Cre (SPOPmut cells) viruses (Augello et al., 2019). The expression of the human *SPOP-F133V* transgene was confirmed by immunoblot. To knockout *Ar* expression, the cells were infected with lentiviruses containing sgNT or sgAr gRNA in pLentiCRISPRv2 plasmids (a kind gift from Dong Zhao, Sawyer’s lab, MSK). Plasmid sequences were verified by Sanger sequencing (Genewiz). Polyclonal populations resistant to puromycin were subjected to immunoblot to verify AR protein expression.

### Immunofluorescence

Organoids were resuspended Cell Recovery Solution (Corning) to melt the matrigel without disrupting the 3D cellular organization. Organoids were pelleted and embedded into fibrinogen and thrombin clots. These were transferred into embedding cassettes and fixed in 10% formalin. The paraffin sections were prepared at The Translational Research Program of the Department of Pathology and Laboratory Medicine at WCM. Immunofluorescent staining was performed as described previously (Augello et al., 2019) by using anti-Ck5, anti-Ck8, and anti-AR primary antibodies (Blattner et al., 2017).

### Pathology review

All murine prostate sections were reviewed by board-certified genitourinary pathologists with expertise in human and murine models of prostate cancer (B.D.R., F.K). All reviews were performed blinded to genotype.

### Immunoblot

Whole-cell protein lysates were prepared after digestion of Matrigel (Corning Cat# 356231) with TrypLE Express Enzyme (1X), phenol red (Gibco). Pelleted cells were washed in PBS and lysed in RIPA buffer (Thermo Fisher Scientific, Cat# PI-89901) supplemented with protease and phosphatase inhibitors (Thermo Fisher Scientific, Cat# 78441). Proteins were quantified by BCA assay (Thermo Fisher Scientific, Cat# 23227) and separated on 4-15% Protein Gels (BioRad, Cat# 4568084). SPOP protein was probed by using in-house made rabbit anti-SPOP (1:1000); for AR and FOXA1, we used following rabbit antibodies: Abcam, ab108341 and Abcam, ab23738. For loading control, we used anti-vinculin (ab129002).

### RNA-seq

Murine organoids were grown for 6 days in media containing 1 nM DHT and then for 24 hr without DHT and EGF. Next, cells were stimulated with 10 nM DHT without EGF for 24 hr and harvested for RNA-seq in biological quadruplicates. Total RNA was extracted, and DNaseI treated by Maxwell 16 LEV simplyRNA Cells Kit (Promega, Cat# AS1270) and Maxwell nucleic-acid extraction instrument (Promega). Nanodrop quantified RNA was checked by Bioanalyzer RNA 6000 Nano Kit (Agilent Technologies). Samples with RNA integrity number > 10 were used for library preparation (TruSeq Stranded mRNA Library Preparation (for Poly-A selection and Stranded RNA-Seq) at the WCM Genomics Core. Libraries were sequenced twice on NextSeq500 (Illumina), High-output mode to generate 75 bp reads at WCM Genomics Core. Obtained FASTQ files were aligned to the mm10 genome by using STAR v2.4.0j (Dobin et al., 2013). The reads were counted with HTSeq (Anders et al., 2015). Differential gene expression was obtained in R with DESeq2 v1.20.0 package (Anders and Huber, 2010). Gene set enrichment analysis (Subramanian et al., 2005) was run in pre-ranked mode to identify enriched signatures in the Molecular Signature Database (MSigDB).

### ATAC-seq

Control and SPOPmut organoids were grown and treated as described for the RNA-seq experiment. The experiment was performed in a biological quadruplicate. The single-cell suspension was obtained after digesting organoids in TryPLE. ATAC-seq was performed on 100,00 nuclei as previously described (Grbesa et al., 2017) with the minor modifications. Nuclei were isolated in buffer containing 10 mM Tris-Cl pH7.5, 10 mM NaCl, 3 mM MgCl_2_, 0.1% Tween (Sigma Millipore, Cat # 11332465001), 0.5% NP-40 (Sigma Millipore, Cat # 11332473001), 0.01% digitonin (Promega, Cat# G9441) and 1x protease inhibitors. Primers that were used for quality checks of the libraries and for assessing mitochondrial DNA contamination are listed in Table S2. Library size was checked with a Bioanalyzer 2100 system (Agilent Technologies) and High Sensitivity DNA Kit (Agilent Technologies, Cat # 5067-4626). Generated DNA fragments were quantified with Qubit dsDNA HS Assay Kit (Invitrogen, Cat # Q3285). Libraries were pooled at equimolar concentration and submitted to WCM Genomics core, where they were paired-end sequenced on HiSeq 4000 system (Illumina) with 50-bp reads. Raw FASTQ files were pre-processed at WCM Genomics core.

### Bioinformatic processing of ATAC-seq data

Quality control checks on raw sequence data were done with FASTQC v.0.11.8 (http://www.bioinformatics.babraham.ac.uk/projects/fastqc/). Paired-end reads were aligned to the mm10 genome using Bowtie2 v.2.3.4.1 in a very sensitive mode. BAM files were sorted and indexed with SAMtools v.1.8. The percentage of mitochondrial reads (Table S1) was calculated with SAMtools using a custom script. All non-unique reads and mitochondrial reads were removed before peak calling. Bigwig files were generated with deepTools v.3.3.1 (Ramirez et al., 2016) bamCoverage tools from reads with min. mapping quality of 10 and using RPGC normalization. Generated bigwig files were visualized with UCSC Genome Browser. Coverage at TSS of RefSeq genes was done with deepTools computeMatrix and plotHeatmap tools (Fig. S1c). The fraction of reads in peaks (FRiP) score was calculated by custom script by using BEDtools2 v.2.27 (Quinlan and Hall, 2010) and reported in Table S1. Data quality was confirmed using Encode guidelines. Peaks were called using Genrich (https://github.com/jsh58/Genrich), as explained on the Harvard

Bioinformatics site (https://informatics.fas.harvard.edu/atac-seq-guidelines.html). As an additional quality control metric, we plotted insert size distribution which confirmed nucleosome-free, mono-and di-nucleosomal distribution of the reads (Fig S1d, S2b). Consensus peaks for each cell type and condition from 4 biological replicas were generated with DiffBind v.3.10 package (Ross-Innes et al., 2012). Differential peak identification (in EdgeR (McCarthy et al., 2012), FDR < 0.01), MA plots, and volcano plots were done in the R software environment v.3.6.0 (The R Foundation) by using DiffBind.

Assignment of ATAC-seq peaks to genes – One Mb regions around the transcription start sites of selected genes were retrieved through the UCSC Table Browser program. Then they were intersected with the bed file containing ATAC-seq peak regions.

Annotation of Genomic Regions – *Cis*-regulatory element analysis of consensus and differential ATAC-seq peaks was performed with the annotatePeaks.pl of Homer v.4.10 software (Heinz et al., 2010). The stacked barplot of enrichment of genomic regions was created in R v3.6.0 with a ggplot2 v3.2.1 package.

Integration with RNA-seq data –was done via Cistrome-GO webserver (Li et al., 2019).

### ChIP-seq experiments and bioinformatic data processing

Control and SPOPmut organoids were grown and treated in biological triplicates, as described in the RNA-seq section. For H3K4me2 ChIP single-cell suspension was crosslinked with 1% methanol-free formaldehyde (Thermo Scientific Pierce, Cat #PI-28906) at 37°C. For AR and FOXA1 ChIP organoids were double fixed (Singh et al., 2019) with 2 mM Di(N-succinimidyl) glutarate (Millipore Signal, Cat #80424-5MG-F) and then 1% formaldehyde. Crosslinking was quenched with 125 mM glycine. The crosslinked organoid pellets were snap-frozen and stored at −80 °C until use. Samples were thawed on ice, lysed in 1% SDS containing buffer supplemented with 1x protease and phosphatase inhibitors, and sonicated for 4 x 10 cycles (30 sec on, 30 sec off) in temperature-controlled Bioruptor 300 (Diagenode). Debris was removed by centrifuging at 14,000 rpm at 4°C. One percent of the supernatant was saved as input, and the rest was added to antibody-coupled Dynabead Protein A (Thermo Fisher Scientific, Cat #10001D) and incubated overnight rocking at 4 °C. Used antibodies were: anti-H3K4me2 ChIP-grade (ab7766), anti-AR PG-21 (Millipore, Cat # 06-680) and anti-FOXA1 ChIP-grade (ab23738). Chromatin was washed on ice with 2x each standard wash buffers (Low-Salt, High-Salt, and LiCl) and finally with TE. Decrosslinking of input and immunoprecipitated chromatin was performed in buffer containing 1% SDS, 0.3M NaCl, and 0.2 mg/mL Proteinase K (Thermo Fisher Scientific, Cat #AM2548) for 16 hr at 65 °C. After 2 hr incubation with RNase A (Thermo Fisher Scientific, Cat #EN0531) decrosslinked material was purified with 2 x AMPure XP beads (Beckman Coulter, Cat #A63881) and eluted in 30 μL of 10 mM Tris-Cl, pH 8. The DNA concentration was measured by Qubit dsDNA HS Assay Kit (Thermo Fisher Scientific, Cat # Q32851). Individual ChIP samples were verified by qPCR (primers are listed in Table S2). Libraries were prepared by NEBNext Ultra II DNA Library Prep (New England Biolabs, Cat #E7645S) by using NEBNext Multiplex Oligos for Illumina (New England Biolabs, Cat #E7335S and #E7500S). Library size and presence of adapters was verified by Bioanalyzer HS DNA chips (Agilent Technologies, Cat# 5067-4626). Libraries passing the quality control were pooled and sequenced at WCM Genomics core for 50 cycles on HiSeq4000 to generate single-end 50-bp reads. Generated FASTQ files were validated for quality using FastQC v.0.11.8 software and processed with ENCODE ChIP-seq pipeline (https://github.com/ENCODE-DCC/chip-seq-pipeline2). All reads were aligned to mouse genome mm10. Briefly, this pipeline generated an HTML-quality report from which we inferred the overall quality of our ChIP-seq. Only the samples passing the ENCODE-specified NSC, RCS, FRiP, and PBC metrics (Landt et al., 2012) (https://www.encodeproject.org/data-standards/chip-seq/) were included in further data analysis (Table S1). All the peaks were called with MACS2(Zhang et al., 2008) at the p-value threshold of 0.05. Differential peak analysis was performed by DiffBind in EdgeR mode with 0.01 FDR cutoff.

### Motif Enrichment Analysis

*De novo* and known motif analysis of ATAC-seq and ChIP-seq consensus or differential peaks was done by findMotifsGenome.pl program within Homer v.4.10 software. For the analysis of H3K4me2 peaks, we used-size given option, while for the rest of the data, we analyzed 200 bp region around the peak summit. To determine motif enrichment between two datasets with similar peak numbers, one served as a background (-bg flag) for the other. For the enhancer-specific motif scanning, peaks localized to promoter and 5’UTR were removed before the analysis. To create motif density histograms the analyzed peaks were centered on the specific motif, and the analysis was run by findMotifsGenome.pl in a histogram mode (-hist with 5 bp bin size) and visualized with GraphPad Prism v8.2.1. Motif analysis of human primary prostate tumors (Stelloo et al., 2018) was done as previously described (Augello et al., 2019) with the only difference that the AR peaks were binned according to the presence or absence of *SPOP* mutation. All the other plots were generated in R with ggplot2 v3.2.1 package.

### Copy number analysis

The copy number alterations of PRAD were downloaded from TCGA portal (https://portal.gdc.cancer.gov/) via gdc-client tool. Fraction of altered cases within *SPOP* and *FOXA1* mutant subclasses were calculated respectively. Segments with log2-ratio >0.3 were defined as genomic amplifications, and log2-ratio < −0.3 were defined as genomic deletions.

### 3D organoid growth assay

Five thousand cells were seeded in 10 μL of matrigel plug (Corning Matrigel Basement Membrane Matrix, Growth Factor Reduced) per well (Corning 96 well assay, flat bottom clear, black, polystyrene plate) in 100 μL of mouse organoid media (Blattner et al., 2017) without EGF and with 0.01 nM DHT. Cells were allowed to form organoids for four days before the bicalutamide treatment. Drug treated cells were assayed for cell viability after four days with CellTiter-Glo 3D assay (Promega), as per manufacturer’s instructions. The experiment was repeated for a total of four biological repeats (technical triplicate for each treatment). Luminescence readings (BioTek Synergy Neo microplate reader with Gen5 software) were analyzed with GraphPad Prism 8.2.1 according to the instructions for the analysis of dose-response data.

### siRNA mediated FOXA1 knockdown

Transient knockdown of *Foxa1* was done by transfecting mouse organoid cells grown on the surface of collagen I coated plates with siRNA (SMARTpool: siGENOME Foxa1 siRNA). For the control, we used the same concentration of scrambled siRNA (siGENOME Non-Targeting Control siRNA Pool). Transfection was done with Lipofectamine RNAiMAX according to the manufacturer’s instructions. After four hours media was replaced and cells were lifted and plated into Matrigel plugs. Bicalutamide treatment and cell viability assay were performed as described above.

### Clinical data analysis

We have developed a novel gene expression signature, classifier SCaPT (Subclass Predictor Based on Transcriptional Data), and decision tree to predict the *SPOP* mutant subclass from RNA gene expression data (Liu et al., 2018). We then classified the *SPOP* mutant subclass in a Decipher retrospective cohort of 1,626 patients (Table S3) (Liu et al., 2018) and evaluated the associations between *SPOP* mutant status and patient outcomes of metastasis mortality with and without ADT treatment based on Kaplan-Meier (KM) analysis.

Prevalence of *SPOP* mutation was analyzed on cBioPortal v3.2.0 platform (Cerami et al., 2012) by using the following datasets (accessed on January 2020): Prostate Adenocarcinoma (TCGA, Cell 2015) (Cancer Genome Atlas Research, 2015), Metastatic Prostate Adenocarcinoma (SU2C/PCF Dream Team) (Abida et al., 2019) and Prostate Cancer (MSKCC) (Abida et al., 2017). For time on ADT analysis, patient’s data from cBioPortal and GitHub (https://github.com/cBioPortal/datahub/tree/master/public/prad_su2c_2019) were combined in R v3.6.0. KM analysis and plots were done in R using packages survival 2.44-1.1 and survminer v0.4.6.

## Acknowledgments

We are indebted to prostate cancer patients and families who contributed to this research. We thank Dr. Dawid Nowak (WCM) and Dr. Jonathan E. Shoag (WCM) for helpful discussion. We thank Dr. Daphne Campigli Di Giammartino (WCM) for helpful advice regarding ChIP-seq experiments. We thank Dr. Mark Rubin for his mentorship and guidance in developing these projects and reagents. We thank Weill Cornell Medicine (WCM) Genomics Core Facility, the Biospecimen and Pathology Core of WCM SPORE in Prostate Cancer and Memorial Sloan Kettering Cancer Center cBioPortal. We are grateful to Dr. Eric Klein, Dr. Bruce J. Trock, Dr. R. Jeffrey Karnes and Dr. Robert B. Den for patient data.

## Supplementary Data

**Fig. S1.**
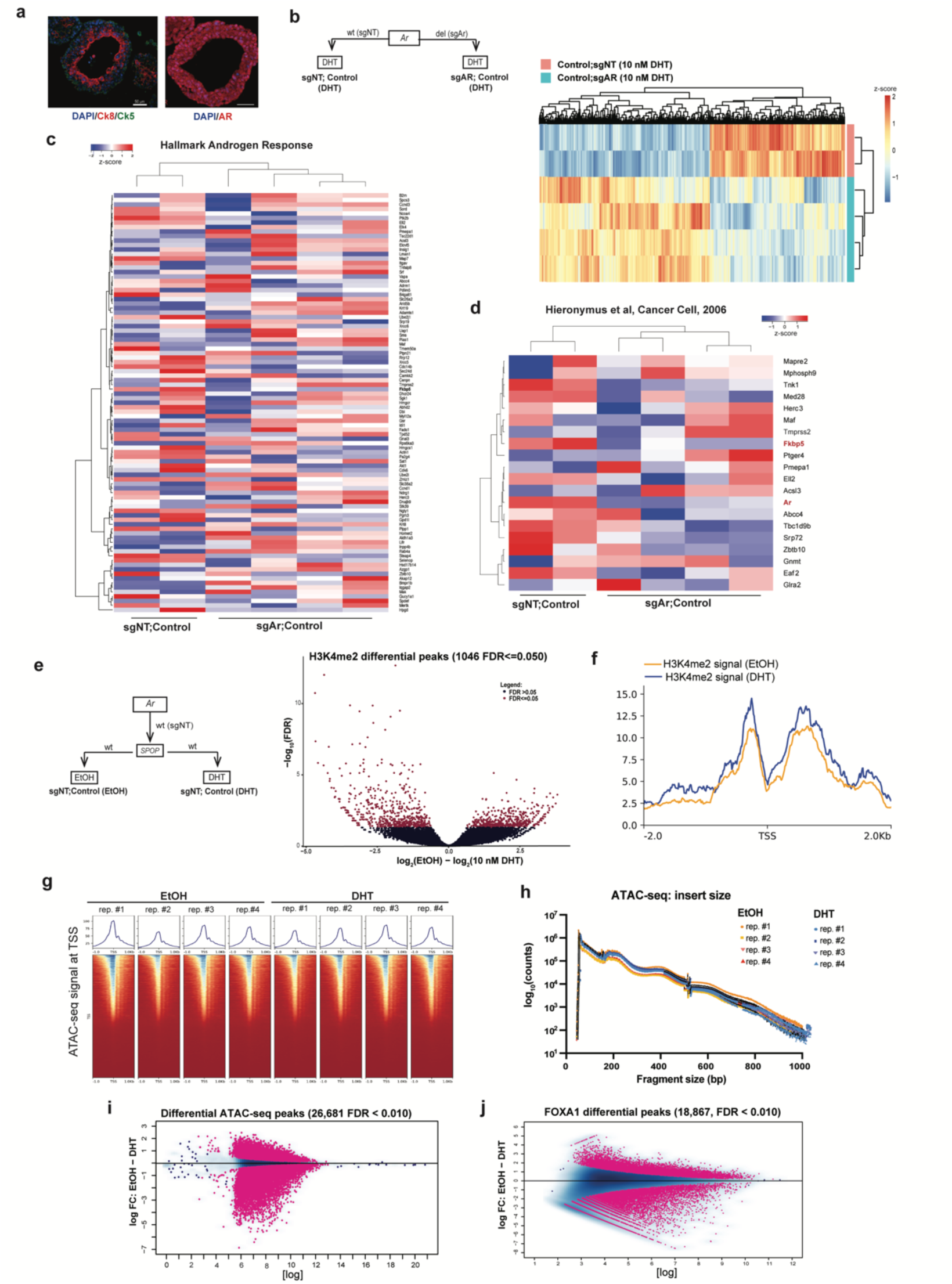
Distinct changes in androgen-induced chromatin and transcriptional landscape of normal murine prostate cells (wildtype SPOP). **(a)** Morphology of murine prostate organoids (left: IF for Ck5 and Ck8) expressing AR (right). **(b)** Schematic representation of RNA-seq experiment and the heatmap showing all the differentially expressed genes. **(c, d)** Response of human AR-target genes, from GSEA **(c)**, and literature (Hieronymus et al., 2006) **(d)**, to Ar downregulation in normal murine prostate organoids. **(e)** Schematics of the ChIP-seq and ATAC-seq experiments (left). Right – Volcano plot showing dynamics H3K4me2 ChIP-seq peaks at FDR 0.05. **(f)** H3K4me2 signal at the promotors of the AR-upregulated genes in Control organoids (shown in Figure 1c). **(g)** ATAC-seq quality metrics - distribution of DNA fragments in generated ATAC-seq libraries. **(h)** ATAC-seq quality metrics – enrichment of ATAC-seq signal at the transcription starts sites (TSS) of housekeeping genes. **(i)** Dynamic ATAC-seq peaks at FDR 0.01. **(j)** Dynamic FOXA1 ChIP-seq peaks at FDR 0.01.

**Fig. S2.**
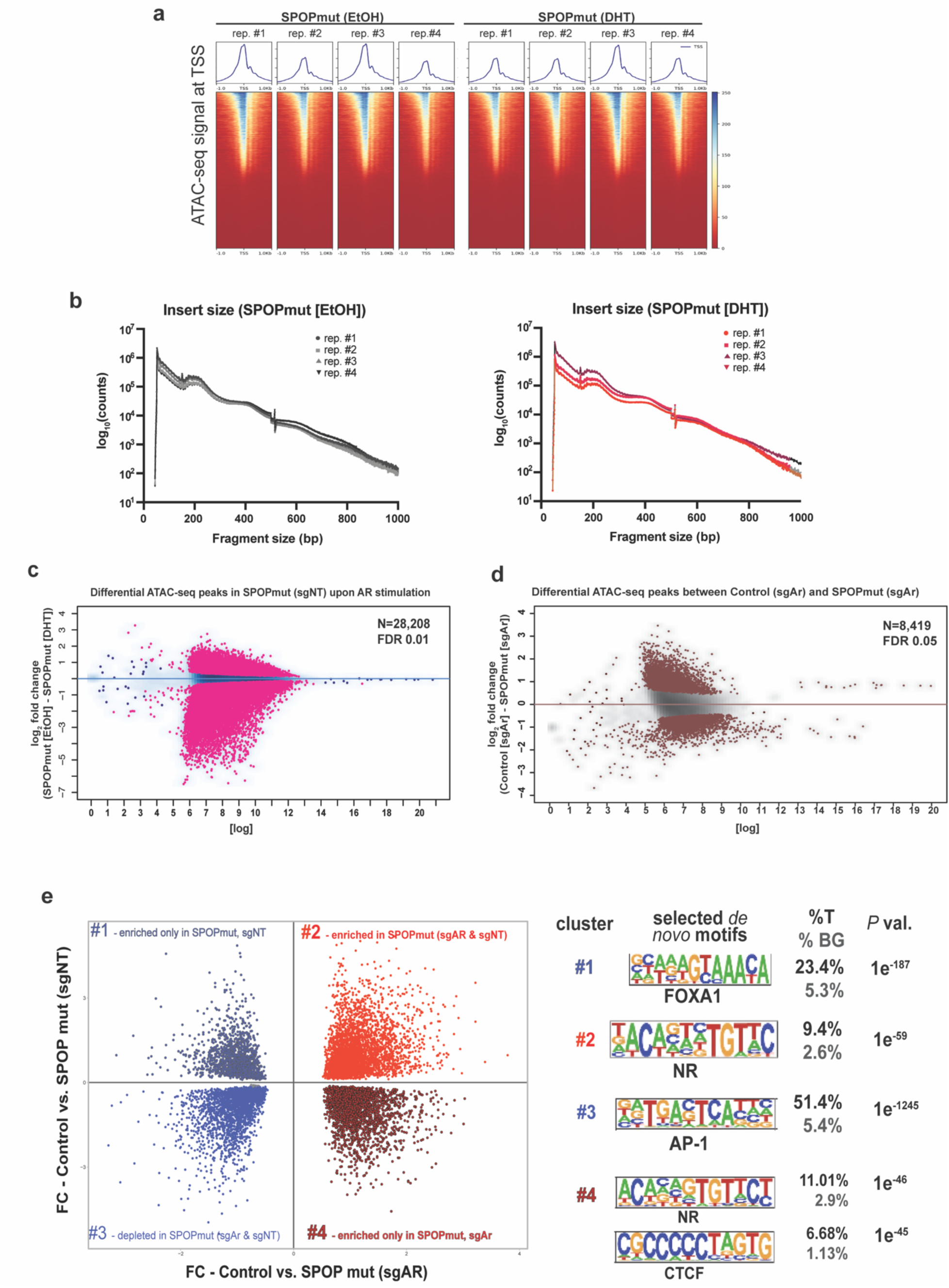
Androgen-dependent and independent chromatin remodeling of murine prostate cells upon introduction of *SPOP* mutation. **(a, b)** Results of Encode quality metrics for ATAC-seq on *SPOP* mutant cells (sgNT) before and after DHT treatment - enrichment at the promoters (i.e. TSS) **(a)** and fragment size distribution **(b)**. **(c)** MA plot – dynamic accessible region upon AR activation. **(d)** MA plot – dynamic regulatory regions induced by *SPOP* mutation when Ar is downregulated. **(e)** Scatter plot – common and unique accessible regions in SPOP mutant cells upon lower Ar expression. Right: results of de novo motif analysis. Black - target regions (%T). Grey - background regions (%BG).

**Fig. S3.**
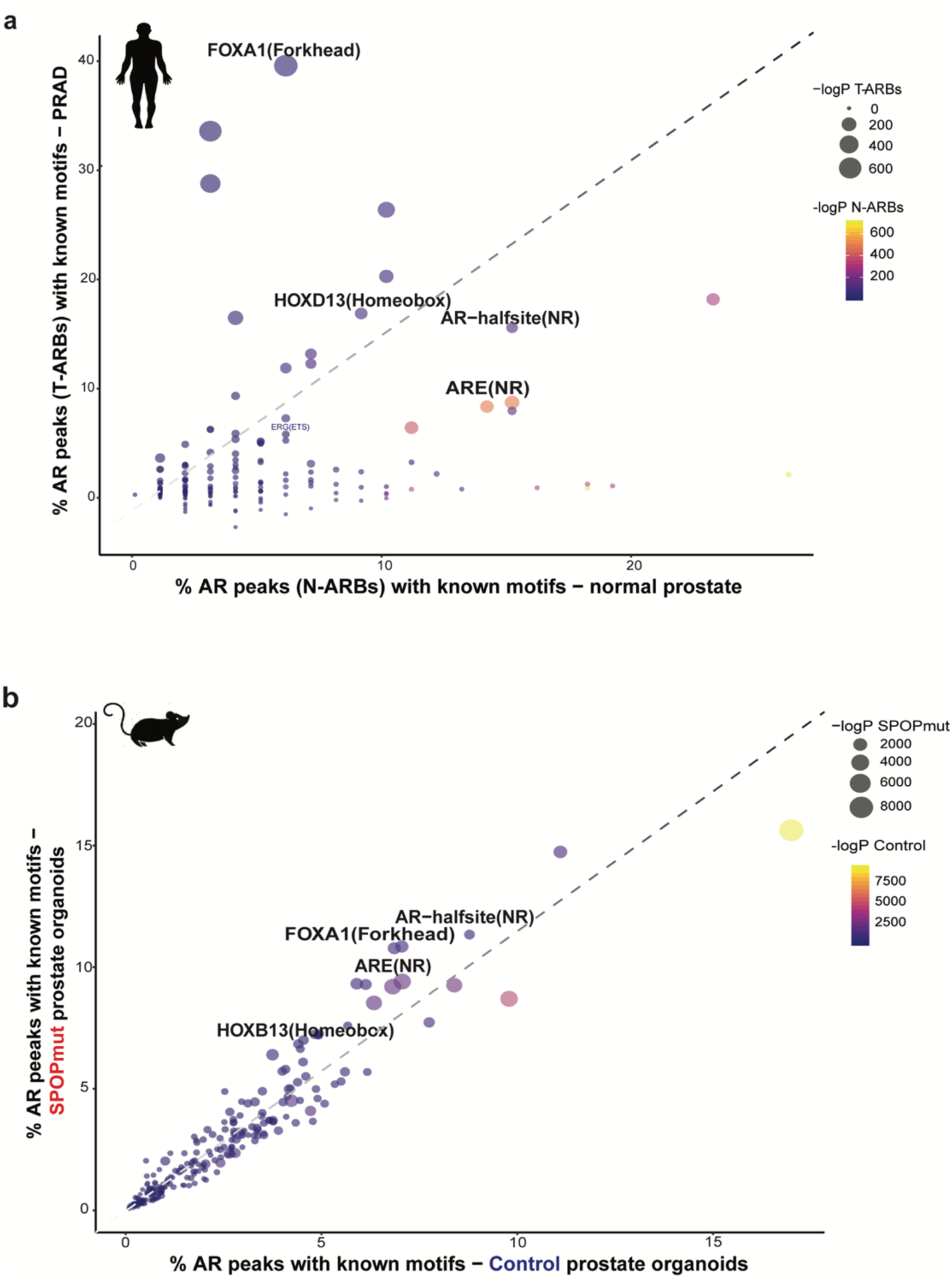
AR cistrome from *SPOP* mutant cells is enriched for FOXA1 and HOXB13 motifs. **(a)** Comparison of the motifs found in AR peaks in normal human prostate tissues and cancers (Pomerantz et al., 2015). **(b)** Motifs found in AR bound regions in before and after introduction of SPOP mutation in normal murine prostate cells.

**Fig. S4.**
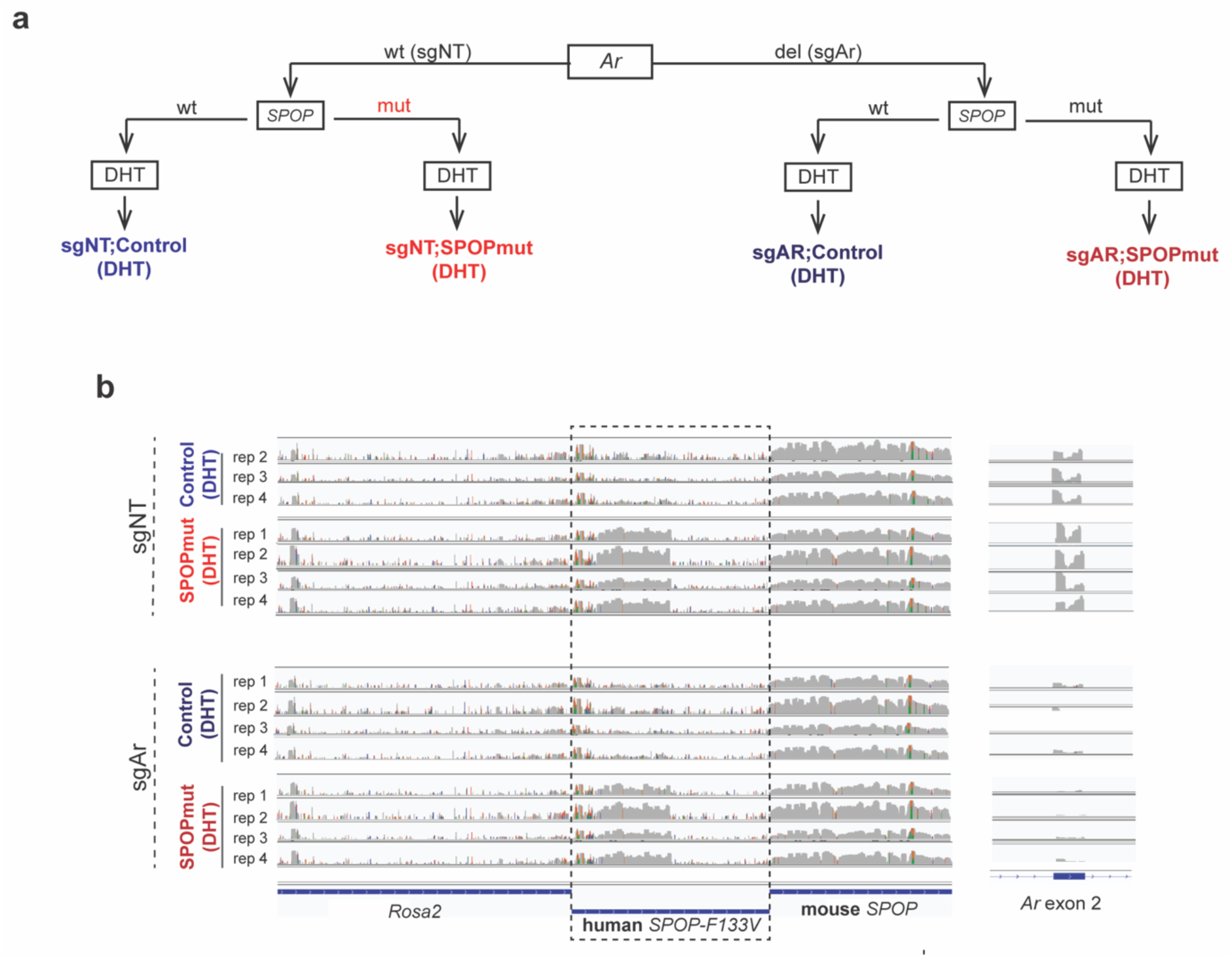
RNA-seq quality control. **(a)** Schematic representation of all the generated RNA-seq libraries. **(b)** Quality controls of the data – reads were aligned to human SPOP sequence (left) and the Ar exon 2 (sgAr target).

**Fig. S5.**
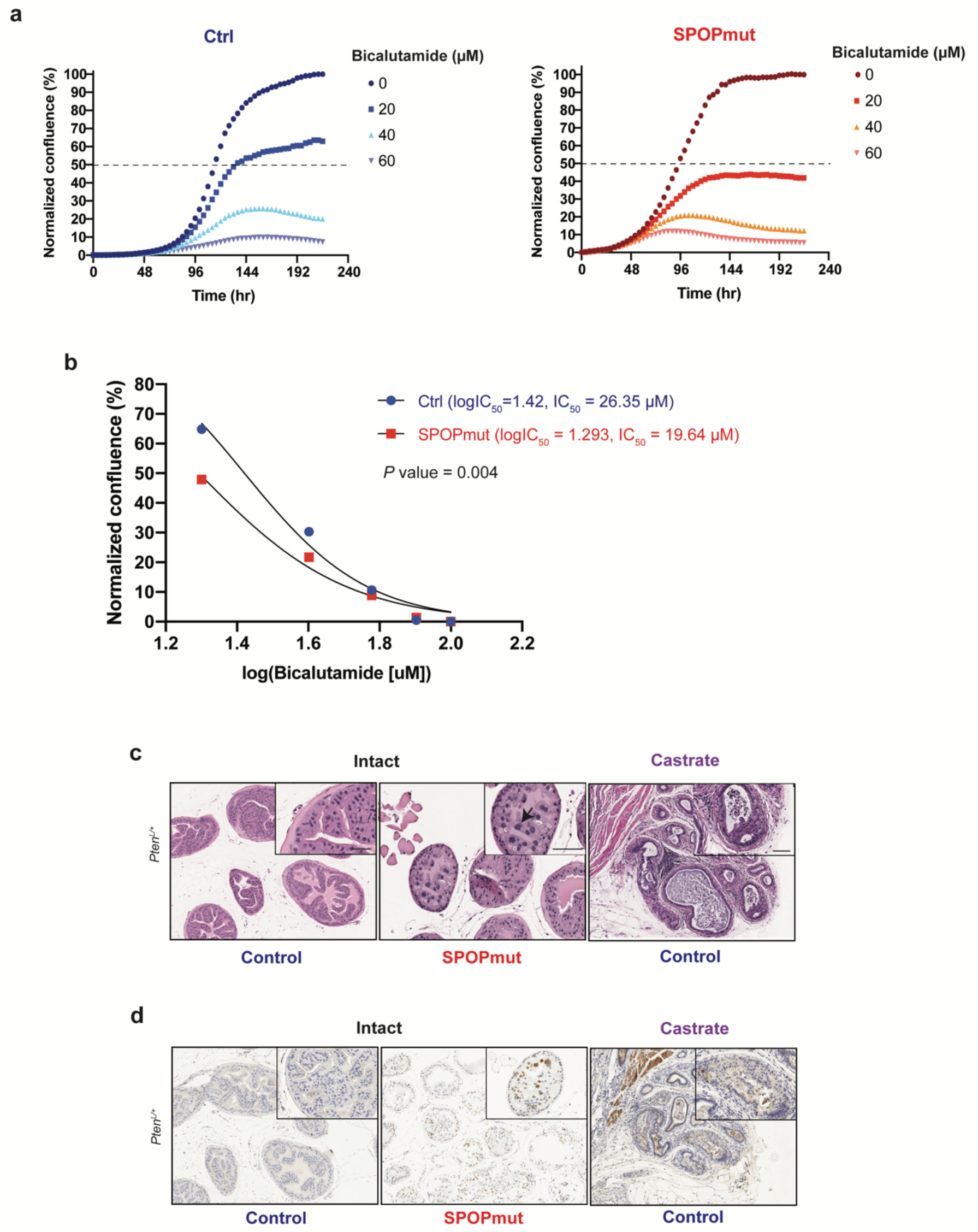
Murine SPOPmut prostate cells show higher sensitivity to anti-androgen treatment. **(a, b)** Slower proliferation of *SPOP*-mutant cells when treated with bicalutamide in 2D (a) and 3D (b). **(c-d)** Representative images showing nuclear atypia and H&E staining.

**Table S3.** Patient’s characteristics from Decipher retrospective cohort used for analysis in Fig.7a.

|  | ***SPOP mutant*** | ***SPOP wild-type*** |
| --- | --- | --- |
| **Cases** | 168 | 1,458 |
| **Median age** (range) | 65 (44-77) | 62 (37-83) |
| **Median PSA** (range) | 8.9 (1.9-87) | 8.6 (0.1-201) |
| **Gleason score** (range) |  |  |
| <=6 | 16 | 176 |
| 3+4 | 30 | 381 |
| 4+3 | 24 | 175 |
| >=8 | 60 | 589 |
| **Pathological tumor stage** |  |  |
| pT2 | 66 | 464 |
| pT3/4 | 82 | 722 |
| **Extracapsular extension** | 88 (53%) | 855 (61%) |
| **Lymp node invasion** | 20 (12%) | 228 (16%) |
| **Seminal vesicle invasion** | 35 (21%) | 452 (32%) |
| **Surgical margin** | 68 (41%) | 703 (49%) |

